# Applying machine learning to identify ionizable lipids for nanoparticle-mediated delivery of mRNA

**DOI:** 10.1101/2023.11.09.565872

**Authors:** Mae M. Lewis, Travis J. Beck, Debadyuti Ghosh

**Author notes:** Corresponding Author; Corresponding Author Address: Division of Molecular Pharmaceutics and Drug Delivery, College of Pharmacy, The University of Texas at Austin, 2409 University Avenue, Austin, TX 78712, USA.

## Abstract

mRNA lipid nanoparticles (LNPs) have a tremendous potential to treat, cure, or prevent many diseases. To identify promising candidates for each application, most studies screen dozens to hundreds of formulations with ionizable lipids synthesized using a single type of chemistry. However, this technique leaves the ionizable lipids synthesized through multi-step chemistries underexplored. This gap in the repertoire of structures is of particular significance because it affects the screening of analogs that are structurally similar to SM-102 and ALC-0315, the ionizable lipids used to formulate the clinically approved mRNA LNP COVID-19 vaccines. Herein, we address this by employing LightGBM, a machine learning algorithm, to reduce the burden of screening these types of ionizable lipids by learning from the breadth and diversity of lipids that have already been tested. We first evaluate the ability of LightGBM to predict LNP potency across heterogeneous chemistries from different studies to achieve an R^2^ of 0.94. After establishing the predictive capacity of the model, we then identify the number of outside carbons in the ionizable lipid as the most important factor contributing to transfection efficiency. From this finding, we subsequently apply the algorithm to predict the effect of formulating nanoluciferase mRNA LNPs using analogs of SM-102 and ALC-0315 with small changes in the number of outside carbons on luciferase activity in HEK293T cells and achieve an R^2^ of 0.83. Importantly, this correlation encompasses novel lipids not included within the database used to train the algorithm. Overall, this study demonstrates the potential of machine learning to accelerate the development of new ionizable lipids by simplifying the screening process.

## Introduction

mRNA therapeutics can be readily tuned to treat a wide range of diseases such as cancer, viral infections, and genetic disorders^1,2^. Lipid nanoparticles (LNPs) have arisen as excellent carriers to facilitate encapsulation and intracellular delivery of therapeutic nucleic acids, as exemplified by their clinical approval for treating hereditary transthyretin-mediated amyloidosis^3^ and for their more recent rapid approval for SARS CoV-2^4,5^. While these trailblazing formulations are effective, the identification of lead candidates is often a time-consuming and costly process. During screening, the formulation efficacy (e.g., transfection efficiency) can be tested by varying each of the traditional four lipid components in LNP formulations: the ionizable lipid, helper lipid, PEG-Lipid, and cholesterol^6^. However, considering structural changes to each component and altering their ratios yields a massive number of possible testable formulations (1×10^10^)^7,8^. To narrow this design space, most studies utilize a fixed ratio of components while focusing their efforts on changing the structure of the ionizable lipid. It is the primary component responsible for complexation with nucleic acids and endosomal escape for intracellular delivery, and many ionizable lipids can be synthesized for extensive screening and testing. As clinically relevant examples, Onpattro and the two COVID-19 vaccines Spikevax and Comirnaty have different ionizable lipids while holding the helper lipid and cholesterol constant and keeping relatively similar component ratios^9^.

Most studies screening for ionizable lipids feature single-step methods of synthesis. For example, thousands of lipidoids have been synthesized through a one-pot three-component reaction with variable polyamine heads, linkers, and alkyl tails^10,11^. Mid-sized libraries with dozens to hundreds of lipids have also been constructed using Michael addition reactions, albeit with more complex amine head groups^12^ and with tails containing heteroatoms^13^. While these methods cover a wealth of structural ground, they leave the structures that require more complex methods of synthesis underexplored. Overall, ionizable lipids synthesized through single-step chemistries have a great deal of diversity from which to learn and apply to these underutilized multi-step chemistries^14,15^. The value of these ionizable lipids requiring more complex chemistries is underscored by the clinically approved mRNA LNPs^16–18^, all of which use ionizable lipids synthesized through multi-step chemistries^6^. However, the lack of comparison amongst different chemistries from different groups makes it challenging to develop a holistic understanding between structure and function of the lipid-based formulations. A method to apply the knowledge gained from other chemistries could be beneficial to simplify the screening process for ionizable lipids and even compare amongst chemistries.

To this end, machine learning (ML) is a powerful approach that can process large, complex data to make models that identify the parameters driving function and predict the behavior of new formulations not even in the dataset. These features highlight the importance of ML in reducing the burden of drug discovery^19^. Within the area of nanomedicine, ensemble algorithms such as gradient boosting, support vector machine, random forest, and artificial neural networks have been applied to analyze structure-function relationships^20–30^. ML has been used to predict behaviors of such as formation of drug nanocrystals^31^, self-emulsifying drug delivery systems^32^, and biomaterials^33,34^, but there are limited reports using this technology to integrate the findings from mRNA LNPs across different chemistries into the development of future formulations^25^. In this study, we capitalize on the power of ML to synthesize the knowledge gained from single-step chemistries and apply it to predict the transfection efficiency of LNPs formulated with ionizable lipids synthesized from multi-step chemistries. We curated a database from publications that formulated four-component LNPs encapsulating reporter luciferase mRNA across studies and labs. We then subsequently applied an ML algorithm, light gradient-boosting machine (LightGBM), to analyze the most important features of ionizable lipids. Leveraging the results from this analysis, we formulated LNPs with analogs of SM-102 and ALC-0315 ionizable lipids with different number of carbons on the outside of the molecule, which was predicted to have the greatest impact on transfection efficiency. Our successful validation of our generated predictions highlights the potential of ML algorithms to accelerate the design and testing process for future ionizable lipid screens.

## Methods

### Extracting data

The dataset was obtained from papers that met the following criteria: (i) formulated four-component mRNA LNPs, (ii) LNPs encapsulated luciferase mRNA, (iii) tested multiple ionizable lipid structures, and (iv) performed testing *in vitro* (Figure 1). Starting with 43 papers that report on luciferase mRNA LNPs as of April 2023, we excluded 29 papers that either (i) added a fifth component (either for stability or targeting purposes), (ii) modified other lipid components (such as helper lipid or cholesterol), or (iii) tested only in vivo. With these exclusions, the final dataset included 2,332 LNPs from 14 different papers across 10 different labs^10–15,35–42^.

**Figure 1.**
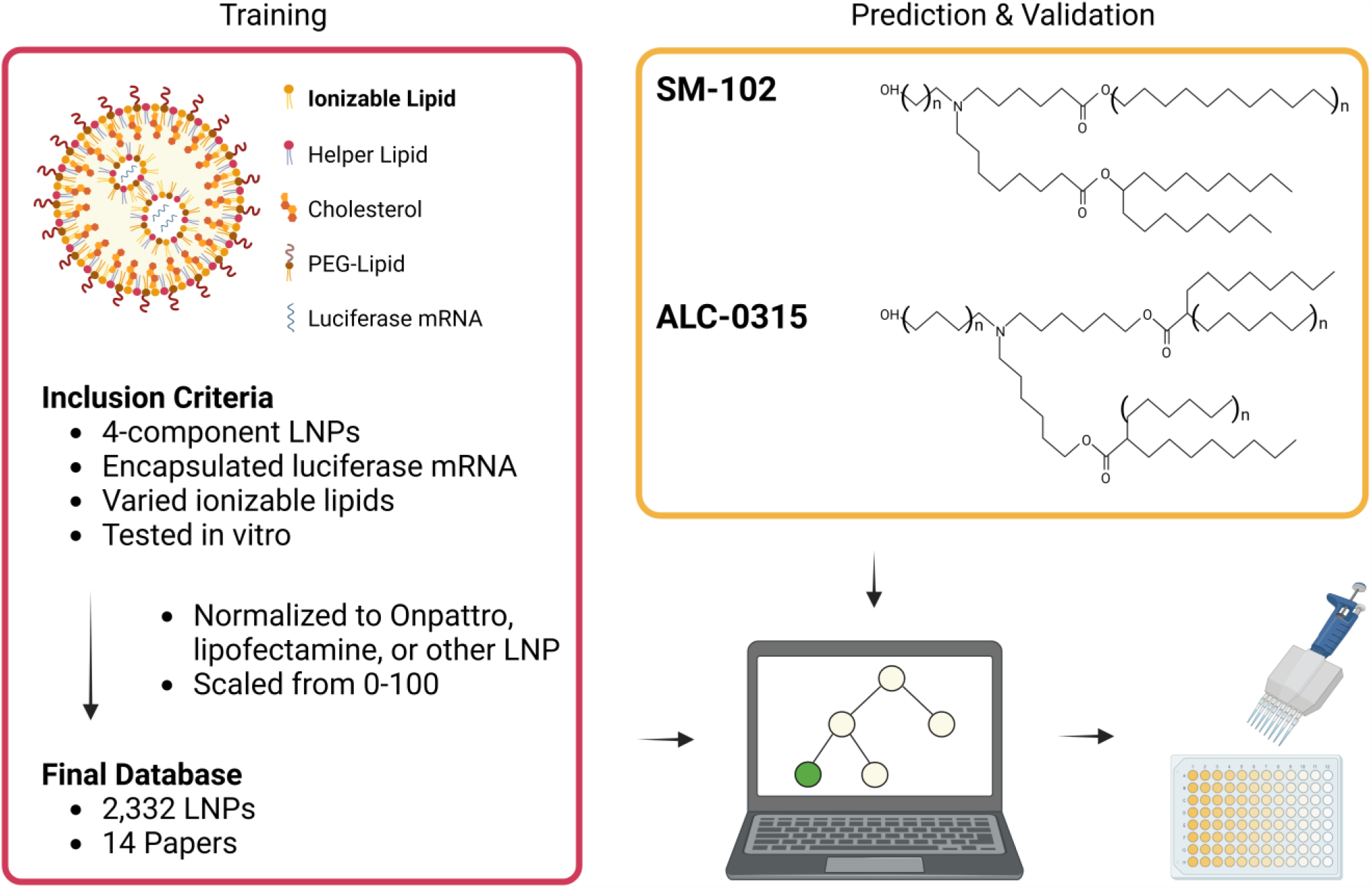
Schematic of ML process. Training LightGBM on the database of four-component luciferase mRNA LNPs. Prediction and validation of mRNA LNPs formulated with analogs of SM-102 and ALC-0315 on HEK293Ts and A549s. Schematic created with BioRender.com.

### Processing data

All mRNA LNPs were formulated with four different types of lipids (ionizable lipid, helper lipid, PEG-Lipid, and cholesterol). These were included in the database as well as their ratios, the lipid to mRNA ratio, and pH of the formulation buffer. Additionally, other experimental conditions (dosage and cell line) were included.

Ionizable lipid chemistry is represented by extended-connectivity fingerprints. These are bit arrays that represent molecular features of a molecule^43^. The presence of “0” in the array means a certain structure is absent while the presence of “1” means the structure is present. To capture all variability within the dataset, the fingerprint radius was set to 5 and the sequence length is set to 4096. Extended connectivity fingerprint arrays were generated by the RDKit package in python.

Ionizable lipids are also represented by structural features on a larger scale that are easily interpretable. We included number of tails, amines, rings, esters, carbons from the amine to the esters, heteroatoms in the tail, double bonds, triple bonds, lipid molecular weight, and a binary category for whether the lipid was branched in the dataset.

Luciferase activity data was extracted by hand. This process involved extracting either raw or normalized luminescence data from bar graphs or from heat maps (Extended Methods). The final database is verified by another individual to ensure accuracy.

As LNPs from each paper were tested in different experimental conditions, we normalized the data by dividing each luciferase reading within one experiment by the highest value and multiplying by 100, putting the final luminescence measurement for each LNP on a scale from 0-100. Additionally, not all experiments used the same controls for comparison. This presented a major hurdle as the ultimate goal was to evaluate the efficacy of LNPs across different chemistries. While some papers used common controls of the Onpattro formulation or lipofectamine, the results of other papers had to be linked based on similarly formulated LNPs (Extended Methods). This scenario affects a majority of the literature (9/14 papers).

### LightGBM

Ensemble modeling ML algorithms have been applied to a wide array of datasets focusing on molecular structure and subsequent effects on performance^25,32,44–48^. Learning from these studies, we assessed the LightGBM algorithm as it is capable of processing relatively small datasets. This algorithm minimizes error by building decision trees sequentially based on a training dataset. In the process of building these trees, LightGBM also trains its parameters more accurately than other algorithms by prioritizing datapoints that have poor predictive power rather than sampling randomly. Within the trees, the leaves will inform the algorithm how to classify the luciferase activity of the LNP given the formulation conditions. After the leaf criteria are determined, the calculated parameters are verified using a smaller subset of data called the testing data. The final luciferase value is an aggregate prediction from each tree that is multiplied by a user-specified learning rate that determines the contribution of each new tree. Overall, gradient boosted decision trees are effective at capturing complex patterns in the data while avoiding underfitting.

The values for parameters such as the number of trees, how many leaves, the rate of sampling, the learning rate, and other parameters can be optimized by the user. We used the sklearn package from Scikit-learn^49^ to employ the LightGBM. We set the learning rate to 0.197, the number of trees to 730, the number of leaves to 700, and the subsample ratio to 0.06. Parameters were optimized using manual search.

A 10-fold cross-validation was performed using ShuffleSplit with a 90:10 train:test split to evaluate the model^25,45^. Sampling was stratified to ensure that each paper was equally represented in both categories. The root mean squared error (RMSE) and R^2^ were calculated to measure model performance.

### mRNA synthesis

mRNA is synthesized as previously described^35^. Briefly, a PCR-amplified NLuc gene block was transcribed using AmpliScribe™ T7-Flash Transcription Kit (Lucigen, ASF-3507) following manufacturer instructions. The mRNA underwent subsequent capping with the cap1 structure using the Vaccinia Capping System (NEB, M2080S) and mRNA Cap 2’-O-methyltransferase (NEB, M0366S). Then, a 3’-poly(A) tail was enzymatically added using the E. Coli Poly (A) Polymerase (NEB, M0276L). Before capping and after polyadenylation, mRNA was purified using RNA Clean & Concentrator-100 (Zymo, R1019). After purification, mRNA was quantified using the Nanodrop One and stored at -80°C until use.

### Lipid selection

The ionizable lipids in the FDA approved COVID-19 vaccines from Moderna and BioNTech-Pfizer, SM-102 and ALC-0315 respectively, as well as their analogs were selected from the ionizable lipid category from BroadPharm (https://broadpharm.com/product-categories/lipid/ionizable-lipid). Lipid selection is limited based on availability. To test the effect of carbons on the outside of SM-102, BP Lipid 114 (9-carbon single chain tail), BP Lipid 113 (7-carbon single chain tail), and BP Lipid 142 (3-carbon head) were selected. To test this for ALC-0315, BP Lipid 226 (2-carbon head), BP Lipid 223 (5-carbon head), and BP 227 (2-carbon head, 8-carbon tails) were selected.

### LNP formulation and characterization

LNPs were formulated using the NanoAssemblr as previously described^35^. Briefly, the lipids combined in 100% EtOH and the NLuc mRNA combined in 50 mM sodium citrate buffer (pH = 3.0) were mixed at a 1:3 organic:aqueous phase using a microfluidic setup. NLuc LNPs were formulated at an mRNA concentration of 10 ng/uL with an *N/P* ratio (ratio of the number of amines in the ionizable lipid to number of phosphates in the mRNA) of 5.67. LNPs were dialyzed against 1x PBS for 4 hours using Slide-A-Lyzer™ G2 Dialysis Cassettes with a 10 kDa molecular weight cut-off (87730, Thermo Fisher). Formulations were stored at 4°C until use.

Encapsulation efficiency was quantified by Ribogreen (R11490, Thermo Fisher Scientific) after dilution to 0.2 ng/uL and DLS using Zetasizer Nano-ZS (Malvern Instruments) after dilution 0.1 ng/uL in 0.1X PBS.

### Luciferase activity in cell culture

HEK293T cells and A549 cells were cultured in Dulbecco’s Modified Eagle’s Medium supplemented with 10% fetal bovine serum and 1% penicillin-streptomycin. Both cell lines were plated at 60,000 cells per well in 100 µL of media in a 96-well plate. After 24 hours, 100 ng of mRNA LNPs in 100 µL of media were added. After a 24-hour incubation, luminescence was read by removing 150 µL of media and adding 50 µL of substrate from the Nano-Glo Luciferase Assay System (N1110, Promega). The mixture was incubated for 3 minutes in a white-walled plate before analysis with a SpectraMax M3 plate reader (Molecular Devices).

### Statistical analysis

Statistical significance for *in vitro* experiments were performed using a double-tailed Student’s t test. All analysis was done in GraphPad Prism (version 8.4.3).

Supplementary figures and extended methods are available in the supplementary information.

## Results

### Machine learning model

The final database of luciferase mRNA LNPs (2,332) is curated following the methods section^10–15,35–42^ (Figure 1). As LNPs encapsulating luciferase mRNA are tested at the beginning stages of screening for many applications, formulations with both low and high transfection are reported. The inclusion of both types of LNP is especially useful for ML algorithms to help delineate in generating models. Notably, we exclude other cargos, LNP formulation types, and LNPs tested only in vivo from our model to avoid contamination of the algorithm with LNPs that may have inherently different performance or poor *in vitro*-*in vivo* correlations. However, differences in experimental technique and methods across labs is still another large hurdle to overcome. By establishing internal controls, we attempt to minimize this interlaboratory variation (Extended Methods).

The final database shows the heterogeneity of ionizable lipid features across publications and labs. Within the ratios of the lipid components, the ionizable lipid ranges from 0.08-0.89, the helper lipid ranges from 0.00-0.50, the PEG-Lipid ranges from 0.00-0.28, and the cholesterol ranges from 0.00-0.66 (Figure 2a). The structural features of the ionizable lipid are also heterogeneous: number of amines (2-9), esters (0-12), rings (0-4), number of tails (1-6), and tail length (5-19) (Figure 2b-f). Additionally, the lipid component ratios in the formulations are highly variable. Only two parameters were largely held constant: (1) DOPE comprised the helper lipid for the vast majority of formulations, and (2) most LNPs are formulated with a lipid to mRNA weight ratio of 10 or 20.

**Figure 2.**
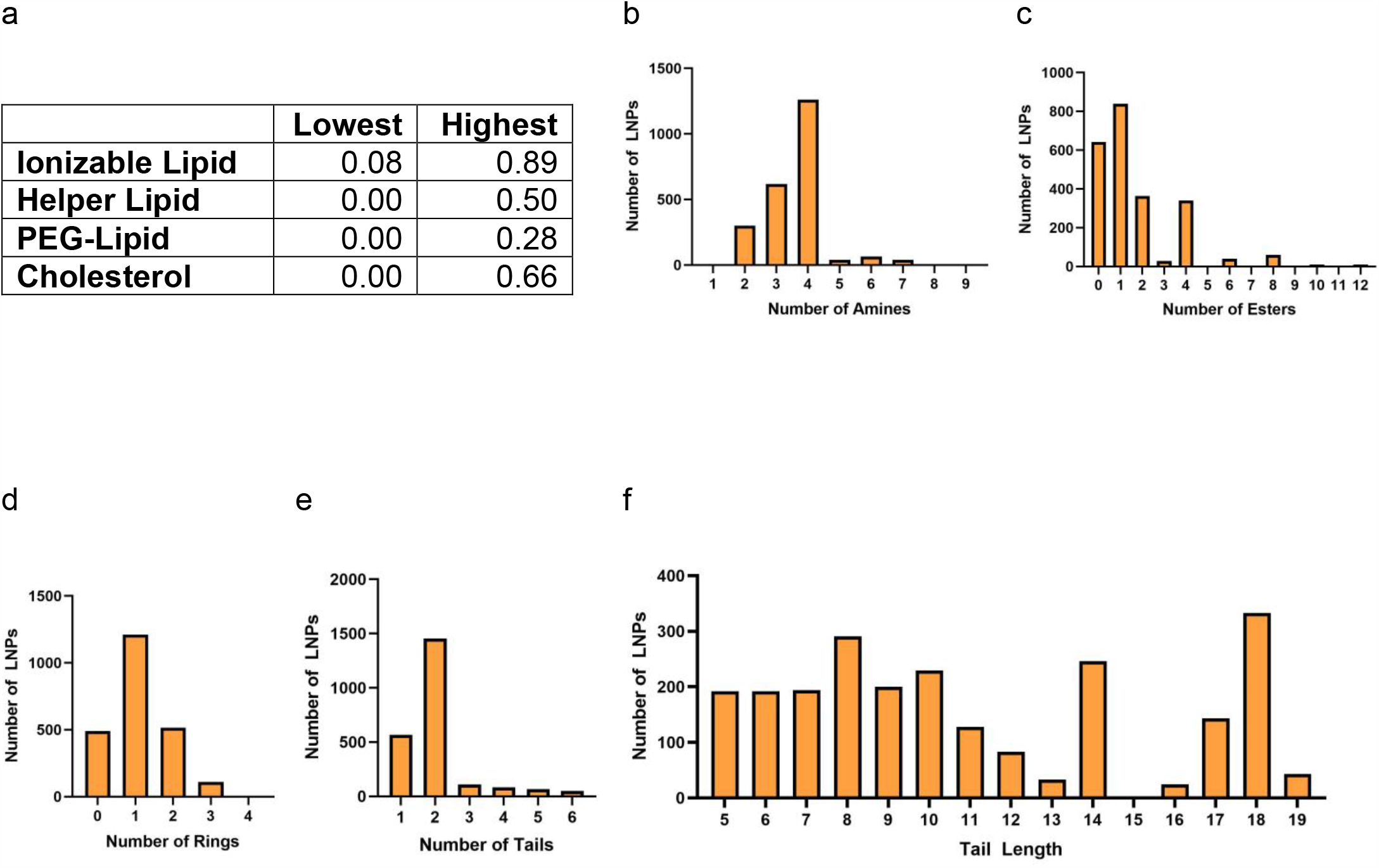
Overview of database statistics. (a) Lowest and highest ratio of each lipid component. (b) Number of amines. (c) Number of esters. (d) Number of rings. (e) Number of tails. (f) Tail length.

After database construction, we explored the relationship between structural features of the ionizable lipid and normalized luminescence. Linear regression coefficients between features such as lipid ratios and structural features of the ionizable lipid poorly predicted the luminescence: the highest R^2^ value was 0.07 (Table S1). We further evaluated ionizable lipid features to identify the highest performing lipids in each category: 5 amines, 10 esters, 0 rings, 5 tails, and 18-carbon tails (Figure S1a-e). As the bulk of the database is comprised of lipidoid structures, these observations reflect the findings of lipid synthesis using epoxide ring opening, Michael addition, and three-component reaction chemistries: a combination of heads with multiple amines and longer alkyl tails yields higher transfection^10,11^. While these categories reveal overarching characteristics about the data, they did not reveal one specific trait that garnered the highest transfection.

We sought to evaluate the capability of an ML algorithm to analyze the features driving transfection efficiency (Figure 1). To prepare the database for this purpose, we represented the ionizable lipid as an extended connectivity fingerprint to capture the chemical heterogeneity more accurately. We then assessed the predictive power of LightGBM. Gradient boosted decision tree algorithms have proven to be a robust predictor of efficacy in other fields of nanomedicine compared to other methods of supervised learning, such as random forest, support vector machine, and logistic regressions^23,25,50^. The R^2^ and RMSE were evaluated with 10-fold cross-fold validation using a 90:10 train:test ratio (Figure 3a) to tune the parameters of the algorithm and evaluate performance. Despite the variability in the dataset, LightGBM was able to achieve an R^2^ of 0.94 ± 0.03 and an RMSE of 1.64 ± 0.72 (Figure 3b).

**Figure 3.**
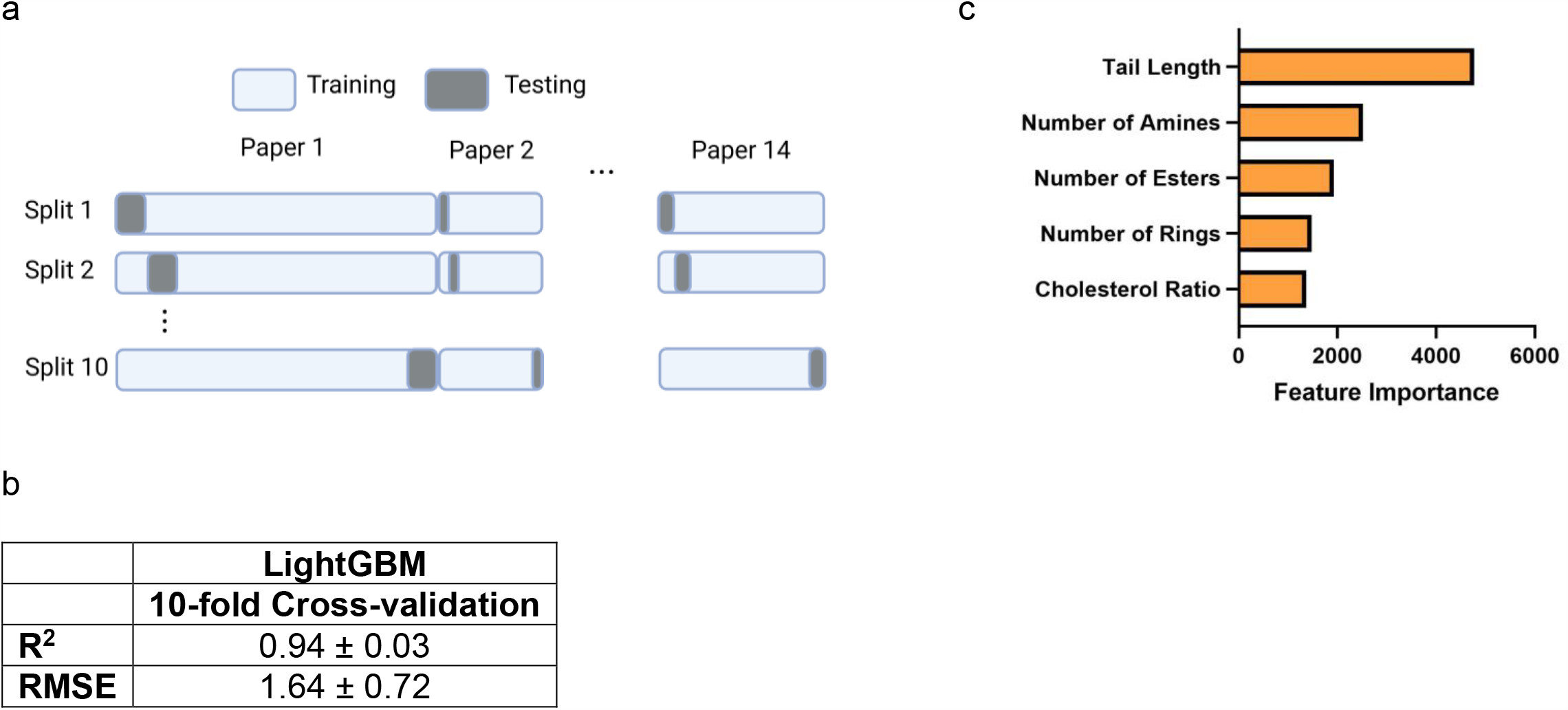
Overview of ML algorithm performance on the database. (a) Schematic of 10-fold cross-validation stratified by paper using a 90:10 train:test ratio. (b) LightGBM statistics (R^2^, RMSE) of 10-fold cross-validation from predicting luminescence. (c) The top five most important features affecting transfection efficiency identified by LightGBM.

### *In vitro* validation of mRNA LNPs

After validating the algorithm and analyzing the database, we employed LightGBM to predict the transfection efficiency of new ionizable analogs based on the feature importance output. LightGBM generates a ranked list of each the effect each LNP formulation parameter has on luminescence through its feature importance function. Based on this analysis, tail length was the most important feature (Figure 3c). As many of the ionizable lipids within this database, we interpreted “tail length” as carbons on the outside of the molecule. Thus, we chose to test variable numbers of carbons measured from the primary amine group and the esters of SM-102 and ALC-0315 (Figure 4a). These analogs were selected from the greatest range of variability available in the BroadPharm catalog. For SM-102, the single chain tail varied from 11 carbons to 9 carbons (BP 114) and to 7 carbons (BP 113) while the head was varied from 2 carbons to 3 carbons (BP 142). For ALC-0315, the carbons from the amine varied from 4 carbons to 5 carbons (BP 223) and to 2 carbons (BP 226). Another lipid, BP 227, was tested to evaluate the effect of multiple changes at once. Compared to ALC-0315, BP 227 is also four-tailed; however, all four-tails have 8 carbons rather than two tails having 8 carbons and the other two having 6 carbons. In addition, BP 227 has a 2-carbon head and the double bonds of the esters are closer to the amine. To evaluate the effects of solely the ionizable lipid, we formulated all LNPs at the same ratio of components as previously validated in our lab (ionizable lipid:DPPC:Cholesterol:DMPE-PEG 2000 45:20:34:1)^35^ to encapsulate nanoluciferase (NLuc) mRNA. Upon formulation, all ionizable lipids showed high encapsulation of mRNA (> 88%) (Figure 4b) and had similar sizes ranging from 62.60 nm to 92.16 nm (Figure 4c). The LNPs were evaluated using the A549 (human bronchial epithelial) and HEK293T (human embryonic kidney) cell lines, which are used in over a third of the database (Figure 4d).

**Figure 4.**
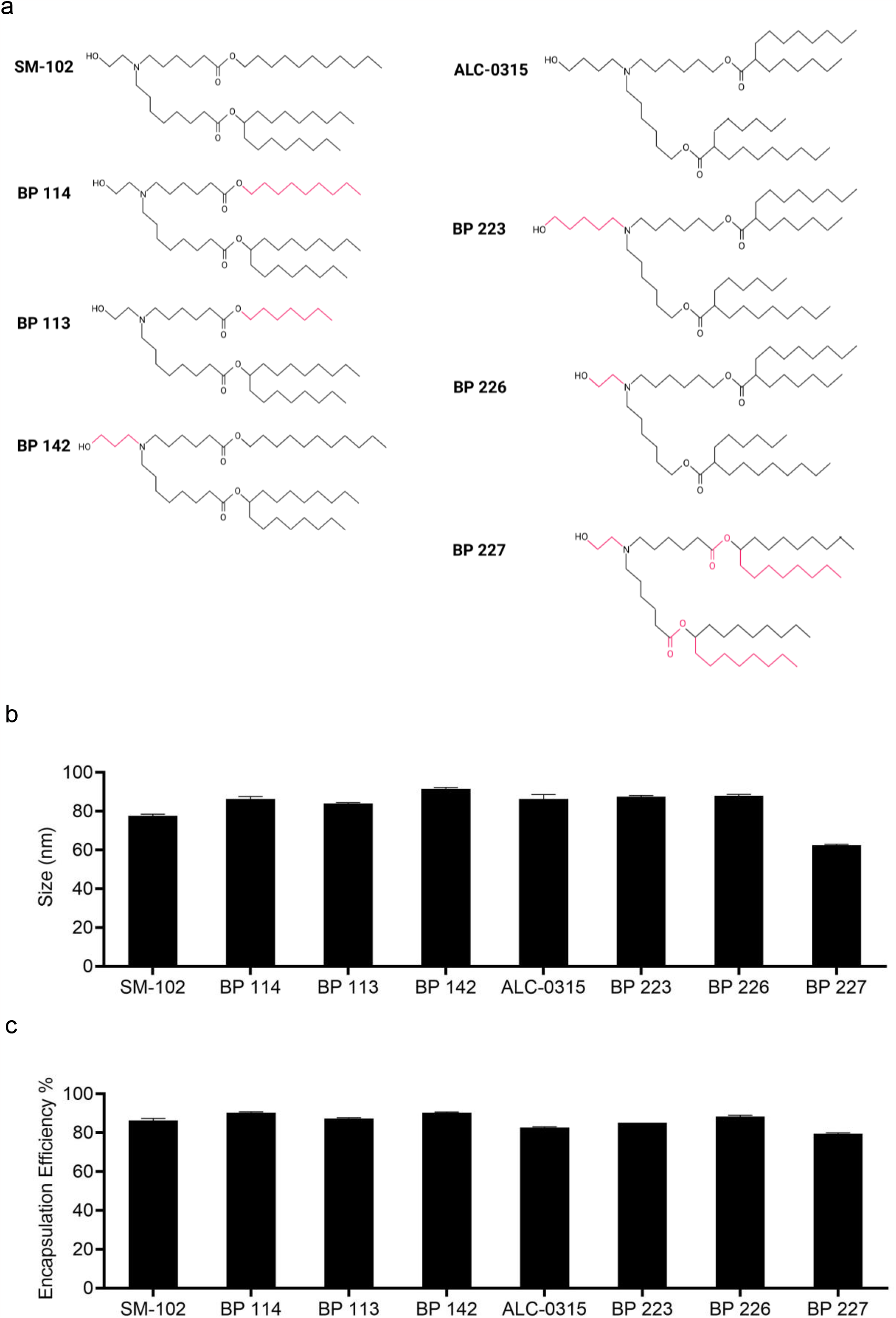
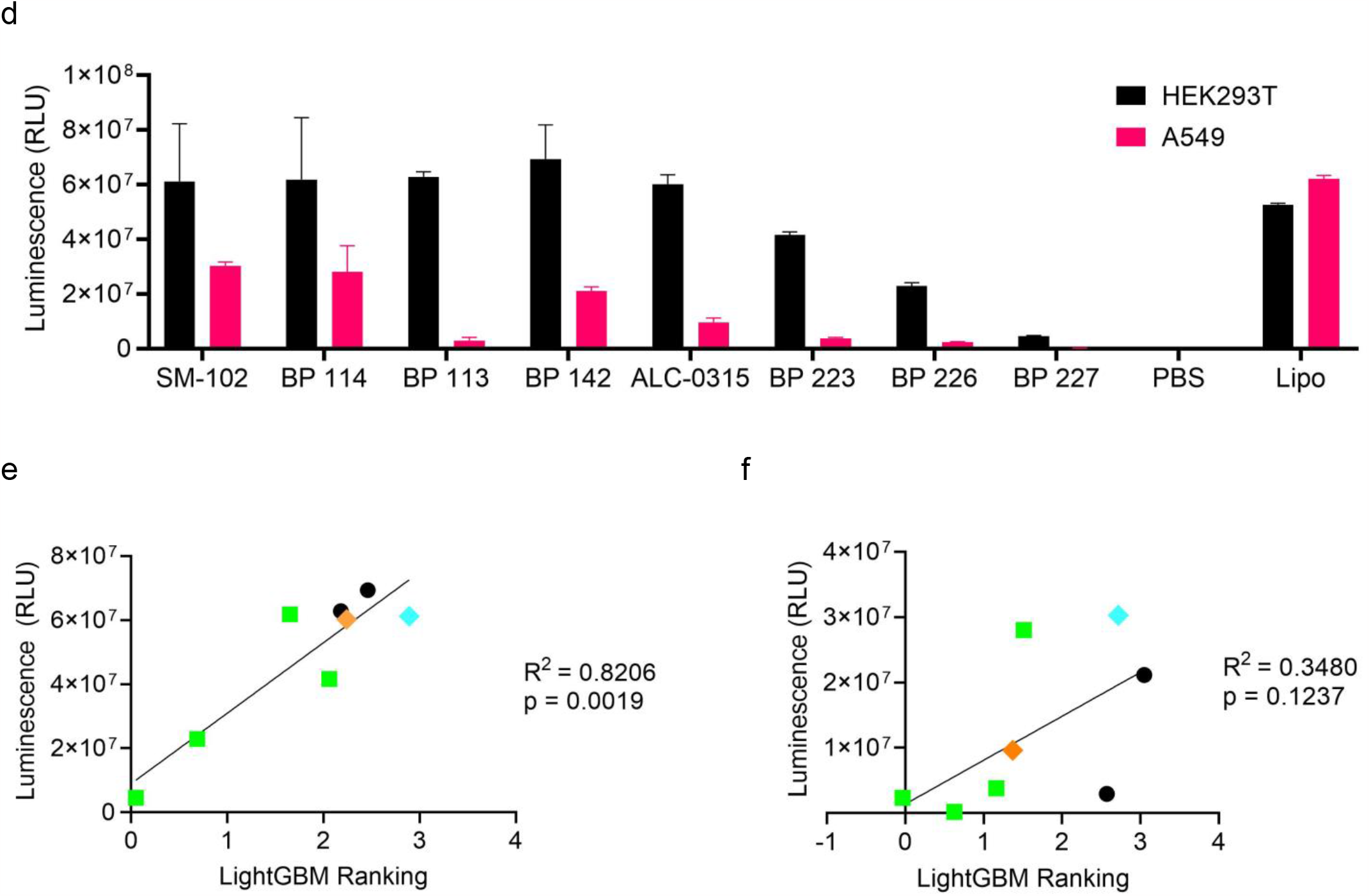
*In vitro* validation of mRNA LNPs formulated with analogs of SM-102 and ALC-0315. (a) Structures of ionizable lipids. Carbons in pink indicate a change from the parent structure. (b) Size (nm) and (c) Encapsulation efficiency of mRNA LNPs upon formulation. (d) Luciferase activity of HEK293T and A549 cells 24 hours after delivery of 100 ng of NLuc mRNA LNPs. R^2^ values of luminescence vs. LightGBM predictions for (e) HEK293Ts and (f) A549s. Light green = ionizable lipids not included within the database. Orange = ALC-0315. Blue = SM-102.

The experimental results of HEK293T cells correlated well with the predictions by the ML algorithm (R^2^ = 0.83) (Figure 4e). While this correlation includes lipids present within the database (SM-102, ALC-0315, BP 142, and BP 113), it also encompasses novel lipids that were not in the database of existing compositions (BP 114, BP 223, BP 226, and BP 227 – highlighted in green). Despite the structural similarity of these novel analogs to their parent lipids, two (BP 114 and BP 223) were predicted to perform similarly compared to ALC-0315 and SM-102 while the other two (BP 226 and BP 227) were predicted to have worse performance. The experimental results validated the algorithm -BP 226 and BP 227 indeed had the lowest transfection efficiency in HEK293Ts.

On the other hand, the luminescence of A549 cells did not correlate well to the model’s predictions (R^2^ = 0.34) (Figure 4f). The main discrepancy lies in the prediction of BP 113, the SM-102 analog with the shortest tail, which performed the worst in the lung epithelial cells. While the decrease in efficiency is substantially worse compared to HEK293T cells, a decrease of this effect size is not recorded in the database. The BP 113 formulated in the same LNP conditions tested on Calu-3s (another lung epithelial cell line) shows a smaller decrease (2.2-fold) than SM-102, compared to the 10.2-fold decrease in A549 cells shown in this study^35^. This could be due to the difference in culturing conditions – the Calu-3 cells were cultured at air-liquid interface, a more physiologically relevant culturing method for lung epithelial cells while the A549s were cultured in submerged conditions.

## Discussion

The intersection of ML and lipid nanoparticle design has enormous potential to reduce the burden of future screens and produce more potent mRNA LNPs. More specifically, ML has the capacity to learn from the heterogeneity of formulations across different chemistries and apply the findings to lipids that are more laborious to synthesize. This is especially beneficial for studying ionizable lipids similar to SM-102 and ALC-0315 -while these lipids are among the first ionizable lipids to be used in chemically approved formulations, only a handful of chemical analogs have been reported due to their multi-step synthesis process^15,17^. To address this problem, we built a database containing thousands of mRNA LNPs with ionizable lipids synthesized spanning different chemistries, analyzed the findings through ML, and applied the resulting algorithm to predict analogs of SM-102 and ALC-0315.

To build the database, we limited our criteria to four-component luciferase mRNA LNPs tested *in vitro*. In this way, we focused our findings primarily on the structure of the ionizable lipid. We first evaluated the capacity of linear correlations to identify drivers for high transfection efficiency. As these failed to isolate a responsible factor, we then evaluated the ability of an ML algorithm, LightGBM, to capture the heterogeneity of the database more accurately. Despite the database containing thousands of lipids across different labs and types of synthesis, LightGBM produced an R^2^ of 0.94 ± 0.03 and an RMSE of 1.64 ± 0.72. These are similar to the results of other nanoparticle studies employing ensemble algorithms to overcome the heterogeneity of small testing sets^25,45,50^, although the RMSE is > 1. More specifically, our study is comparable to a publication employing LightGBM to analyze the IgG titer of 325 samples of four-component mRNA LNP across multiple studies which achieves an R^2^ of 0.87 ± 0.06 and an RMSE of 0.41 ± 0.09 with 10-fold cross-validation^25^. While the applications of our results diverge (Wang et al. validates the predictive power of LightGBM on lipids included within the database whereas we use feature importance to inform testing of novel lipids not included within the database), both of our studies underscore the utility of gradient boosting decision trees to analyze compiled databases of mRNA LNPs.

The feature importance generated by LightGBM revealed that the structural features of the ionizable lipids had the greatest impact on mRNA delivery. More specifically, the tail length, or number of carbon chains on the outside of the molecule, was the most impactful feature. Therefore, we evaluated the effect of changing both the head and tail length of the ionizable lipids ALC-0315 and SM-102 on transfection efficiency. We formulated the mRNA LNPs based on our previous work^35^ to validate the findings in cell lines commonly found in the database (HEK293T and A549). For HEK293Ts, the algorithm predicted the experimental data well (R^2^ = 0.83). In this cell line, SM-102 analogs performed similarly to the parent structure. The highest performer was BP-142 (3-carbon head), although its luminescence was not significantly greater than that of SM-102 in this cell line. This lipid, however, has resulted in significantly higher luminescence compared to SM-102 using another cell line (Calu-3s) and equivalent LNP formulation conditions^35^. The lowest performing LNP, BP 227, was an analog of ALC-0315 that varied the number of carbons in the head group and tail groups while flipping the position of the oxygens in the esters. While minor changes in carbon numbers in the other analogs of SM-102 and ALC-0315 were mostly comparable to their respective structures, this larger shift away from the parent structure produced the greatest reduction in delivery. Therefore, the importance of SM-102 and ALC-0315 is not a function of one particular structure; any major changes can abrogate the function. This is in line with the study of other lipid libraries that demonstrate the dramatic decrease on potency by changes in even one carbon^10,11,14^. One library of helper lipids studies the underlying mechanism behind this decrease through endosomal escape. In a screen of 572 helper lipids, a three alkyl-tailed lipid containing 10-12 carbons in combination with a small zwitterion head were more effective at promoting the formation of an inverse hexagonal phase H^II^ at endosomal pH^51^. Tail lengths outside of this range were significantly less effective. Based on our study and previous screens, the tail length is of critical importance across chemistries and is therefore a great candidate for further improvement by ML and by future mechanism studies.

While the prediction model worked well for HEK293T cells, the linear regression for A549s was not significant and had an R^2^ of 0.34. The main discrepancy between the experimental data and the algorithm is BP 113 (7-carbon tail). While this lipid performed similarly to SM-102 in HEK293Ts, it performed significantly worse in A549s. The inability of LightGBM to predict this result may be due to the types of ionizable lipids in the database tested on HEK293Ts vs A549s. In the database, HEK293Ts are used in 3 publications, all of which use MC3 or analogs of MC3^37–39^. However, A549s are only used in one publication which tests analogs of C12-200, a different class of lipid with distinct chemical structures compared to MC3, SM-102, and ALC-0315^11^. We hypothesize that molecular modeling of the interactions between the LNPs and its binding receptors on the cell surface as well as its interaction with the endosomal membrane could shed light on the difference in performance.

While this study pools data from other chemistries for the benefit of testing SM-102 and ALC-0315 analogs, the algorithm could be improved with a larger dataset, better experimental standardization, and more comprehensive reporting methods for LNP formulation conditions. In addition, this study only considers luminescence data (i.e., luciferase activity). Other factors, such as cargo and immunogenicity, have an effect on the performance of LNPs as well. Our exclusion of certain criteria localizes our work to the structure of the ionizable lipid. However, the screens of other lipid components not included in this study, such as β-sitosterol as a cholesterol analog or 9A1P9 as a helper lipid, have demonstrated superior transfection. These components, however, have only been tested with a handful of ionizable lipids^36,51,52^. To leverage these findings to establish more potent LNPs, these components should be tested in conjunction with a wider range of established excipients. In this way, an ML algorithm can better predict beneficial interactions. Overall, future generations of mRNA LNPs will greatly benefit from these proposed studies and from internal standard controls, such as Onpattro or lipofectamine, to compare between chemistries.

## Conclusions

We established a method of compiling a database of luciferase mRNA LNPs across different labs and demonstrated the efficacy of an ML algorithm, LightGBM, in synthesizing this data to predict the transfection efficiency of mRNA LNPs formulated with analogs of SM-102 and ALC-0315. While further exploration of the difference between cell types could improve the model, these findings highlight the advantages of using ML to analyze heterogeneous chemistries to benefit the future design of ionizable lipids.

## Acknowledgements

We would like to acknowledge the Cystic Fibrosis Foundation (GHOSH 19XX0) and Emily’s Entourage for support. We would also like to thank Yuting Pan and Melissa R. Soto for insightful discussions on experimental design. We thank Dr. Pengyu Ren for providing guidance and expertise on the manuscript. Additionally, we would like to thank Dr. Anna Blakney (University of British Columbia) for providing raw data for the database. Some schematics were created with BioRender.com.

## CRediT authorship contribution statement

**Mae M. Lewis:** Conceptualization (lead), Methodology (lead), Software (equal), Validation (equal), Formal analysis (lead), Investigation (lead), Data Curation (supporting), Writing – Original Draft (lead). **Travis J. Beck:** Software (equal), Validation (equal), Data Curation (lead), Writing – Review & Editing (supporting). **Debadyuti Ghosh:** Funding acquisition (lead), Writing – Review & Editing (supporting).

## Notes

### Competing Interest Statement

The authors have declared no competing interest.

## References

1. Gupta A, Andresen JL, Manan RS, Langer R. Nucleic acid delivery for therapeutic applications. Adv Drug Deliv Rev. 2021;178:113834. doi:10.1016/j.addr.2021.113834

2. Hou X, Zaks T, Langer R, Dong Y. Lipid nanoparticles for mRNA delivery. Nat Rev Mater. 2021;6(12):1078–1094. doi:10.1038/s41578-021-00358-0

3. Akinc A, Maier MA, Manoharan M, Fitzgerald K, Jayaraman M, Barros S, Ansell S, Du X, Hope MJ, Madden TD, Mui BL, Semple SC, Tam YK, Ciufolini M, Witzigmann D, Kulkarni JA, van der Meel R, Cullis PR. The Onpattro story and the clinical translation of nanomedicines containing nucleic acid-based drugs. Nat Nanotechnol. 2019;14(12):1084–1087. doi:10.1038/s41565-019-0591-y

4. Baden LR, El Sahly HM, Essink B, Kotloff K, Frey S, Novak R, Diemert D, Spector SA, Rouphael N, Creech CB, McGettigan J, Khetan S, Segall N, Solis J, Brosz A, Fierro C, Schwartz H, Neuzil K, Corey L, Gilbert P, Janes H, Follmann D, Marovich M, Mascola J, Polakowski L, Ledgerwood J, Graham BS, Bennett H, Pajon R, Knightly C, Leav B, Deng W, Zhou H, Han S, Ivarsson M, Miller J, Zaks T. Efficacy and safety of the mRNA-1273 SARS-CoV-2 vaccine. N Engl J Med. 2021;384(5):403–416. doi:10.1056/NEJMoa2035389

5. Polack FP, Thomas SJ, Kitchin N, Absalon J, Gurtman A, Lockhart S, Perez JL, Pérez Marc G, Moreira ED, Zerbini C, Bailey R, Swanson KA, Roychoudhury S, Koury K, Li P, Kalina WV, Cooper D, Frenck RW, Hammitt LL, Türeci Ö, Nell H, Schaefer A, Ünal S, Tresnan DB, Mather S, Dormitzer PR, Şahin U, Jansen KU, Gruber WC. Safety and efficacy of the BNT162b2 mRNA Covid-19 vaccine. N Engl J Med. 2020;383(27):2603–2615. doi:10.1056/NEJMoa2034577

6. Han X, Zhang H, Butowska K, Swingle KL, Alameh MG, Weissman D, Mitchell MJ. An ionizable lipid toolbox for RNA delivery. Nat Commun. 2021;12:7233. doi:10.1038/s41467-021-27493-0

7. Lokugamage MP, Sago CD, Dahlman JE. Testing thousands of nanoparticles in vivo using DNA barcodes. Curr Opin Biomed Eng. 2018;7:1–8. doi:10.1016/j.cobme.2018.08.001

8. Lokugamage MP, Vanover D, Beyersdorf J, Hatit MZC, Rotolo L, Echeverri ES, Peck HE, Ni H, Yoon JK, Kim Y, Santangelo PJ, Dahlman JE. Optimization of lipid nanoparticles for the delivery of nebulized therapeutic mRNA to the lungs. Nat Biomed Eng. 2021;5(9):1059–1068. doi:10.1038/s41551-021-00786-x

9. Schoenmaker L, Witzigmann D, Kulkarni JA, Verbeke R, Kersten G, Jiskoot W, Crommelin DJA. mRNA-lipid nanoparticle COVID-19 vaccines: Structure and stability. Int J Pharm. 2021;601:120586. doi:10.1016/j.ijpharm.2021.120586

10. Miao L, Li L, Huang Y, Delcassian D, Chahal J, Han J, Shi Y, Sadtler K, Gao W, Lin J, Doloff JC, Langer R, Anderson DG. Delivery of mRNA vaccines with heterocyclic lipids increases anti-tumor efficacy by STING-mediated immune cell activation. Nat Biotechnol. 2019;37(10):1174–1185. doi:10.1038/s41587-019-0247-3

11. Li B, Manan RS, Liang SQ, Gordon A, Jiang A, Varley A, Gao G, Langer R, Xue W, Anderson D. Combinatorial design of nanoparticles for pulmonary mRNA delivery and genome editing. Nat Biotechnol. Published online March 30, 2023:1-6. doi:10.1038/s41587-023-01679-x

12. Billingsley MM, Singh N, Ravikumar P, Zhang R, June CH, Mitchell MJ. Ionizable lipid nanoparticle-mediated mRNA delivery for human CAR T cell engineering. Nano Lett. 2020;20(3):1578–1589. doi:10.1021/acs.nanolett.9b04246

13. Zhao X, Chen J, Qiu M, Li Y, Glass Z, Xu Q. Imidazole-based synthetic lipidoids for in vivo mRNA delivery into primary T lymphocytes. Angew Chem, Int Ed Engl. 2020;59(45):20083–20089. doi:10.1002/anie.202008082

14. Li B, Luo X, Deng B, Wang J, McComb DW, Shi Y, Gaensler KML, Tan X, Dunn AL, Kerlin BA, Dong Y. An orthogonal array optimization of lipid-like nanoparticles for mRNA delivery in vivo. Nano Lett. 2015;15(12):8099–8107. doi:10.1021/acs.nanolett.5b03528

15. Naidu GS, Yong SB, Ramishetti S, Rampado R, Sharma P, Ezra A, Goldsmith M, Hazan-Halevy I, Chatterjee S, Aitha A, Peer D. A combinatorial library of lipid nanoparticles for cell type-specific mRNA delivery. Adv Sci. 2023;10(19):2301929. doi:10.1002/advs.202301929

16. Jayaraman M, Ansell SM, Mui BL, Tam YK, Chen J, Du X, Butler D, Eltepu L, Matsuda S, Narayanannair JK, Rajeev KG, Hafez IM, Akinc A, Maier MA, Tracy MA, Cullis PR, Madden TD, Manoharan M, Hope MJ. Maximizing the potency of siRNA lipid nanoparticles for hepatic gene silencing In vivo. Angew Chem, Int Ed Engl. 2012;51(34):8529–8533. doi:10.1002/anie.201203263

17. Sabnis S, Kumarasinghe ES, Salerno T, Mihai C, Ketova T, Senn JJ, Lynn A, Bulychev A, McFadyen I, Chan J, Almarsson Ö, Stanton MG, Benenato KE. A novel amino lipid series for mRNA delivery: Improved endosomal escape and sustained pharmacology and safety in non-human primates. Mol Ther. 2018;26(6):1509–1519. doi:10.1016/j.ymthe.2018.03.010

18. Hassett KJ, Benenato KE, Jacquinet E, Lee A, Woods A, Yuzhakov O, Himansu S, Deterling J, Geilich BM, Ketova T, Mihai C, Lynn A, McFadyen I, Moore MJ, Senn JJ, Stanton MG, Almarsson Ö, Ciaramella G, Brito LA. Optimization of lipid nanoparticles for intramuscular administration of mRNA vaccines. Mol Ther Nucleic Acids. 2019;15:1–11. doi:10.1016/j.omtn.2019.01.013

19. Meyer TA, Ramirez C, Tamasi MJ, Gormley AJ. A user’s guide to machine learning for polymeric biomaterials. ACS Polym Au. 2023;3(2):141–157. doi:10.1021/acspolymersau.2c00037

20. Findlay MR, Freitas DN, Mobed-Miremadi M, Wheeler KE. Machine learning provides predictive analysis into silver nanoparticle protein corona formation from physicochemical properties. Environ Sci: Nano. 2018;5(1):64–71. doi:10.1039/C7EN00466D

21. Lazarovits J, Sindhwani S, Tavares AJ, Zhang Y, Song F, Audet J, Krieger JR, Syed AM, Stordy B, Chan WCW. Supervised learning and mass spectrometry predicts the in vivo fate of nanomaterials. ACS Nano. 2019;13(7):8023–8034. doi:10.1021/acsnano.9b02774

22. Ban Z, Yuan P, Yu F, Peng T, Zhou Q, Hu X. Machine learning predicts the functional composition of the protein corona and the cellular recognition of nanoparticles. Proc Natl Acad Sci U S A. 2020;117(19):10492–10499. doi:10.1073/pnas.1919755117

23. Yamankurt G, Berns EJ, Xue A, Lee A, Bagheri N, Mrksich M, Mirkin CA. Exploration of the nanomedicine-design space with high-throughput screening and machine learning. Nat Biomed Eng. 2019;3(4):318–327. doi:10.1038/s41551-019-0351-1

24. Nademi Y, Tang T, Uludağ H. Modeling uptake of polyethylenimine/short interfering RNA nanoparticles in breast cancer cells using machine learning. Adv Nanobiomed Res. 2021;1(10):2000106. doi:10.1002/anbr.202000106

25. Wang W, Feng S, Ye Z, Gao H, Lin J, Ouyang D. Prediction of lipid nanoparticles for mRNA vaccines by the machine learning algorithm. Acta Pharm Sin B. 2022;12(6):2950–2962. doi:10.1016/j.apsb.2021.11.021

26. Hasanzadeh A, Hamblin MR, Kiani J, Noori H, Hardie JM, Karimi M, Shafiee H. Could artificial intelligence revolutionize the development of nanovectors for gene therapy and mRNA vaccines? Nano Today. 2022;47:101665. doi:10.1016/j.nantod.2022.101665

27. Han R, Ye Z, Zhang Y, Cheng Y, Zheng Y, Ouyang D. Predicting liposome formulations by the integrated machine learning and molecular modeling approaches. Asian J Pharm Sci. 2023;18(3):100811. doi:10.1016/j.ajps.2023.100811

28. Xu Y, Ma S, Cui H, Chen J, Xu S, Wang K, Varley A, Lu RXZ, Wang B, Li B. AGILE platform: a deep learning-powered approach to accelerate LNP development for mRNA delivery. Published online June 2, 2023:2023.06.01.543345. doi:10.1101/2023.06.01.543345

29. Ding DY, Zhang Y, Jia Y, Sun J. Machine learning-guided lipid nanoparticle design for mRNA delivery. Published online August 2, 2023. doi:10.48550/arXiv.2308.01402

30. Bao Z, Bufton J, Hickman RJ, Aspuru-Guzik A, Bannigan P, Allen C. Revolutionizing drug formulation development: The increasing impact of machine learning. Adv Drug Deliv Rev. 2023;202:115108. doi:10.1016/j.addr.2023.115108

31. He Y, Ye Z, Liu X, Wei Z, Qiu F, Li HF, Zheng Y, Ouyang D. Can machine learning predict drug nanocrystals? J Control Release. 2020;322:274–285. doi:10.1016/j.jconrel.2020.03.043

32. Gao H, Jia H, Dong J, Yang X, Li H, Ouyang D. Integrated in silico formulation design of self-emulsifying drug delivery systems. Acta Pharm Sin B. 2021;11(11):3585–3594. doi:10.1016/j.apsb.2021.04.017

33. Li F, Han J, Cao T, Lam W, Fan B, Tang W, Chen S, Fok KL, Li L. Design of self-assembly dipeptide hydrogels and machine learning via their chemical features. Proc Natl Acad Sci U S A. 2019;116(23):11259–11264. doi:10.1073/pnas.1903376116

34. Tamasi MJ, Patel RA, Borca CH, Kosuri S, Mugnier H, Upadhya R, Murthy NS, Webb MA, Gormley AJ. Machine learning on a robotic platform for the design of polymer–protein hybrids. Adv Mater. 2022;34(30):e2201809. doi:10.1002/adma.202201809

35. Lewis MM, Soto MR, Maier EY, Wulfe SD, Bakheet S, Obregon H, Ghosh D. Optimization of ionizable lipids for aerosolizable mRNA lipid nanoparticles. Bioeng Transl Med. Published online August 21, 2023:e10580. doi:10.1002/btm2.10580

36. Patel S, Ashwanikumar N, Robinson E, Xia Y, Mihai C, Griffith JP, Hou S, Esposito AA, Ketova T, Welsher K, Joyal JL, Almarsson Ö, Sahay G. Naturally-occurring cholesterol analogues in lipid nanoparticles induce polymorphic shape and enhance intracellular delivery of mRNA. Nat Commun. 2020;11(1):983. doi:10.1038/s41467-020-14527-2

37. Zhang H, Leal J, Soto MR, Smyth HDC, Ghosh D. Aerosolizable lipid nanoparticles for pulmonary delivery of mRNA through design of experiments. Pharmaceutics. 2020;12(11):1042. doi:10.3390/pharmaceutics12111042

38. Carrasco MJ, Alishetty S, Alameh MG, Said H, Wright L, Paige M, Soliman O, Weissman D, Cleveland TE, Grishaev A, Buschmann MD. Ionization and structural properties of mRNA lipid nanoparticles influence expression in intramuscular and intravascular administration. Commun Biol. 2021;4(1):1–15. doi:10.1038/s42003-021-02441-2

39. Li Z, Zhang XQ, Ho W, Li F, Gao M, Bai X, Xu X. Enzyme-catalyzed one-step synthesis of ionizable cationic lipids for lipid nanoparticle-based mRNA COVID-19 vaccines. ACS Nano. 2022;16(11):18936–18950. doi:10.1021/acsnano.2c07822

40. Young RE, Nelson KM, Hofbauer SI, Vijayakumar T, Alameh MG, Weissman D, Papachristou C, Gleghorn JP, Riley RS. Lipid nanoparticle composition drives mRNA delivery to the placenta. Published online December 22, 2022:2022.12.22.521490. doi:10.1101/2022.12.22.521490

41. Ly HH, Daniel S, Soriano SKV, Kis Z, Blakney AK. Optimization of lipid nanoparticles for saRNA expression and cellular activation using a design-of-experiment approach. Mol Pharmaceutics. 2022;19(6):1892–1905. doi:10.1021/acs.molpharmaceut.2c00032

42. Swingle KL, Safford HC, Geisler HC, Hamilton AG, Thatte AS, Billingsley MM, Joseph RA, Mrksich K, Padilla MS, Ghalsasi AA, Alameh MG, Weissman D, Mitchell MJ. Ionizable lipid nanoparticles for in vivo mRNA delivery to the placenta during pregnancy. J Am Chem Soc. 2023;145(8):4691–4706. doi:10.1021/jacs.2c12893

43. Rogers D, Hahn M. Extended-connectivity fingerprints. J Chem Inf Model. 2010;50(5):742–754. doi:10.1021/ci100050t

44. Jabed MA, Kim Y, Yarbrough C, Harman-Ware AE, Olstad J, Seiser R, Paeper C, Starace AK, Kim S. A machine learning model for predicting composition of catalytic coprocessing products from molecular beam mass spectra. ACS Sustainable Chem Eng. 2023;11(32):11912–11923. doi:10.1021/acssuschemeng.3c01821

45. Yu F, Wei C, Deng P, Peng T, Hu X. Deep exploration of random forest model boosts the interpretability of machine learning studies of complicated immune responses and lung burden of nanoparticles. Sci Adv. 2021;7(22):eabf4130. doi:10.1126/sciadv.abf4130

46. Maharjan R, Hada S, Lee JE, Han HK, Kim KH, Seo HJ, Foged C, Jeong SH. Comparative study of lipid nanoparticle-based mRNA vaccine bioprocess with machine learning and combinatorial artificial neural network-design of experiment approach. Int J Pharm. 2023;640:123012. doi:10.1016/j.ijpharm.2023.123012

47. Han R, Xiong H, Ye Z, Yang Y, Huang T, Jing Q, Lu J, Pan H, Ren F, Ouyang D. Predicting physical stability of solid dispersions by machine learning techniques. J Control Release. 2019;311-312:16–25. doi:10.1016/j.jconrel.2019.08.030

48. Schletz D, Breidung M, Fery A. Validating and utilizing machine learning methods to investigate the impacts of synthesis parameters in gold nanoparticle synthesis. J Phys Chem C. 2023;127(2):1117–1125. doi:10.1021/acs.jpcc.2c07578

49. Pedregosa F, Varoquaux G, Gramfort A, Michel V, Thirion B, Grisel O, Blondel M, Prettenhofer P, Weiss R, Dubourg V, Vanderplas J, Passos A, Cournapeau D, Brucher M, Perrot M, Duchesnay É. Scikit-learn: machine learning in Python. J Mach Learn Res. 2011;12(85):2825–2830.

50. Gao H, Ye Z, Dong J, Gao H, Yu H, Li H, Ouyang D. Predicting drug/phospholipid complexation by the lightGBM method. Chem Phys Lett. 2020;747:137354. doi:10.1016/j.cplett.2020.137354

51. Liu S, Cheng Q, Wei T, Yu X, Johnson LT, Farbiak L, Siegwart DJ. Membrane-destabilizing ionizable phospholipids for organ-selective mRNA delivery and CRISPR–Cas gene editing. Nat Mater. 2021;20(5):701–710. doi:10.1038/s41563-020-00886-0

52. Zeng Y, Escalona-Rayo O, Knol R, Kros A, Slütter B. Lipid nanoparticle-based mRNA candidates elicit potent T cell responses. Biomater Sci. 2023;11(3):964–974. doi:10.1039/D2BM01581A

