## Supplementary Information for "Applying machine learning to identify ionizable lipids for nanoparticle-mediated delivery of mRNA"

**Supplementary Figures**

|  | **R^2^** |
| --- | --- |
| **Number of Tails** | 0.03 |
| **Number of Amines** | 0.01 |
| **Number of Rings** | 0.01 |
| **Number of Esters** | 0.03 |
| **Amine to Ester** | 0.07 |
| **Heteroatoms in Tail** | 0.01 |
| **Tail Length** | 0.03 |
| **Number of Double Bonds** | 0.01 |
| **Number of Triple Bonds** | 0.00 |
| **Lipid Molecular Weight** | 0.07 |

**Table S1. Linear correlation between select features and measured luminescence.**

| 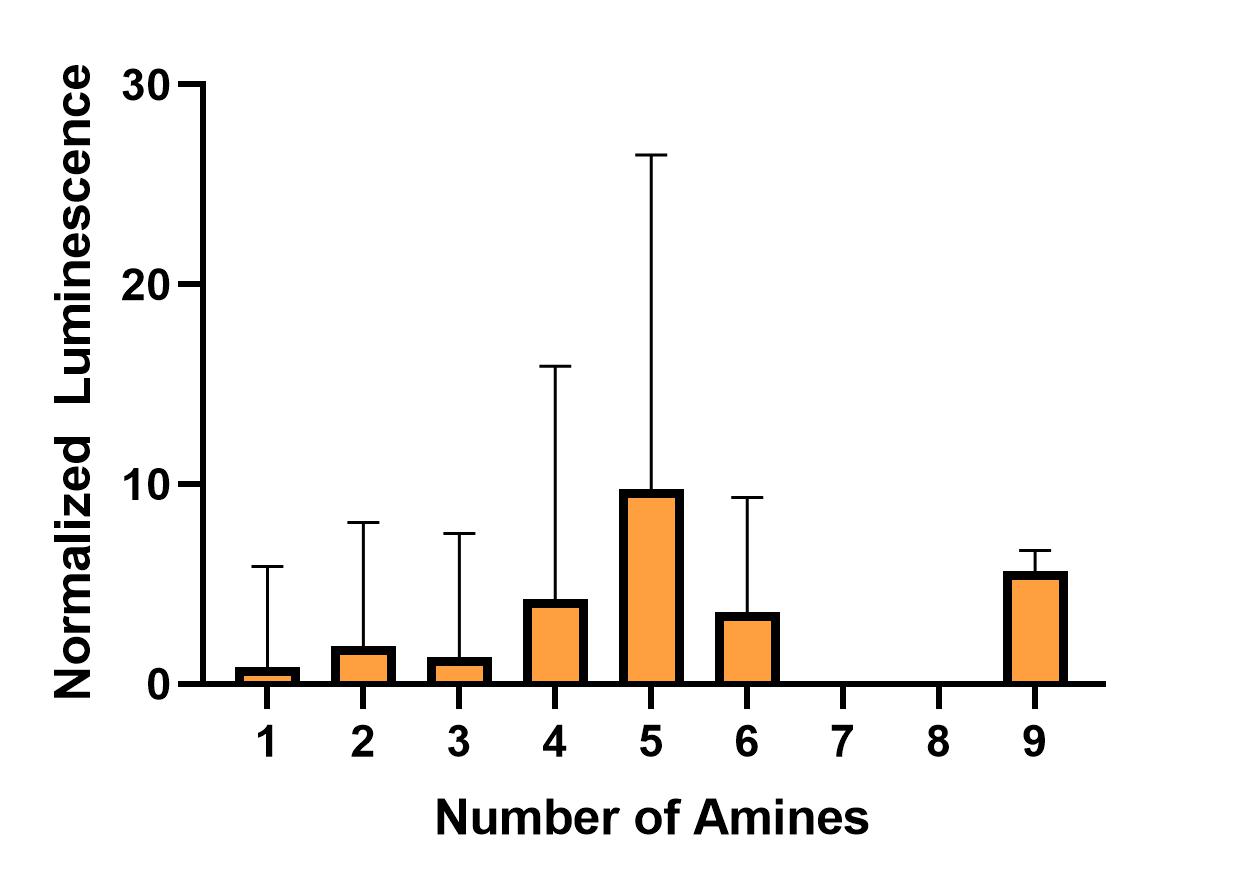a | 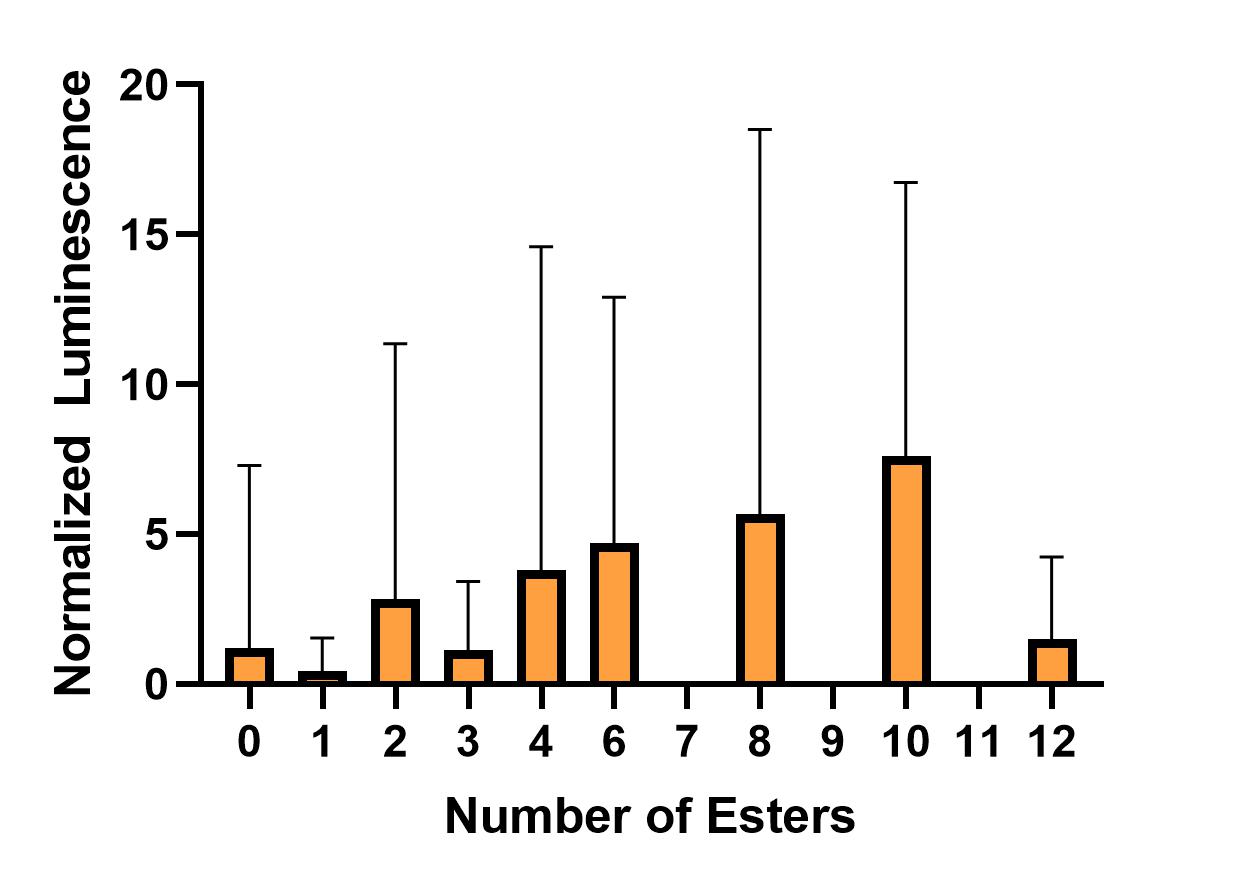b | 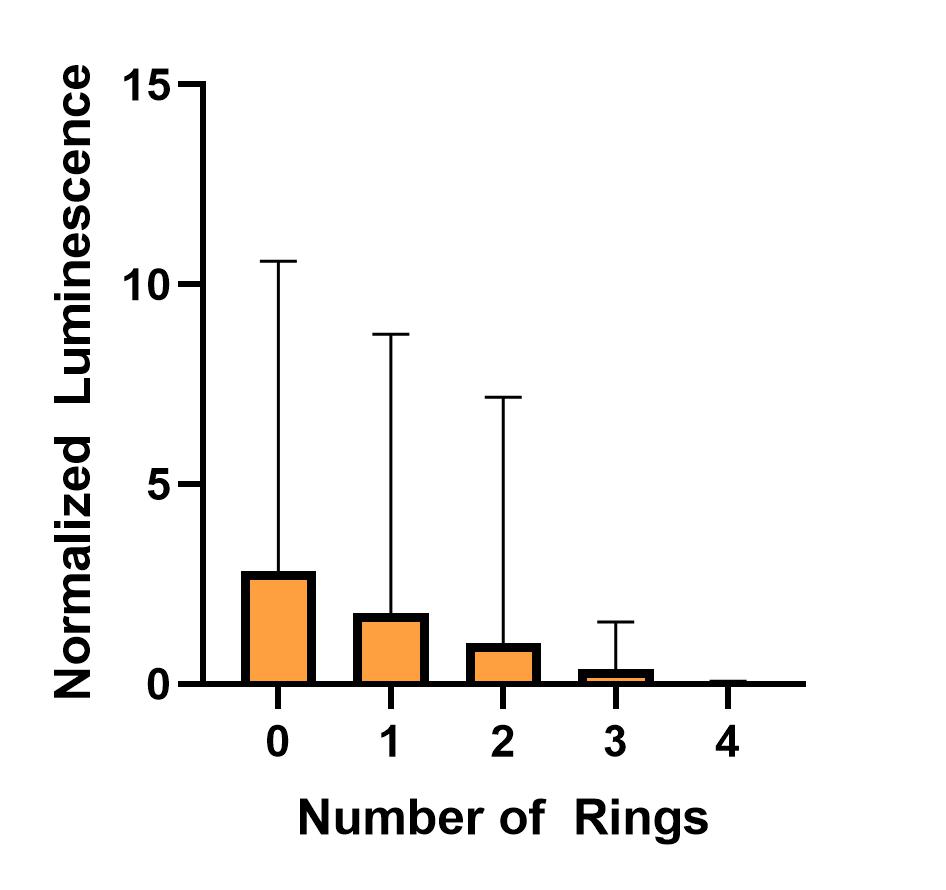c |
| --- | --- | --- |
| 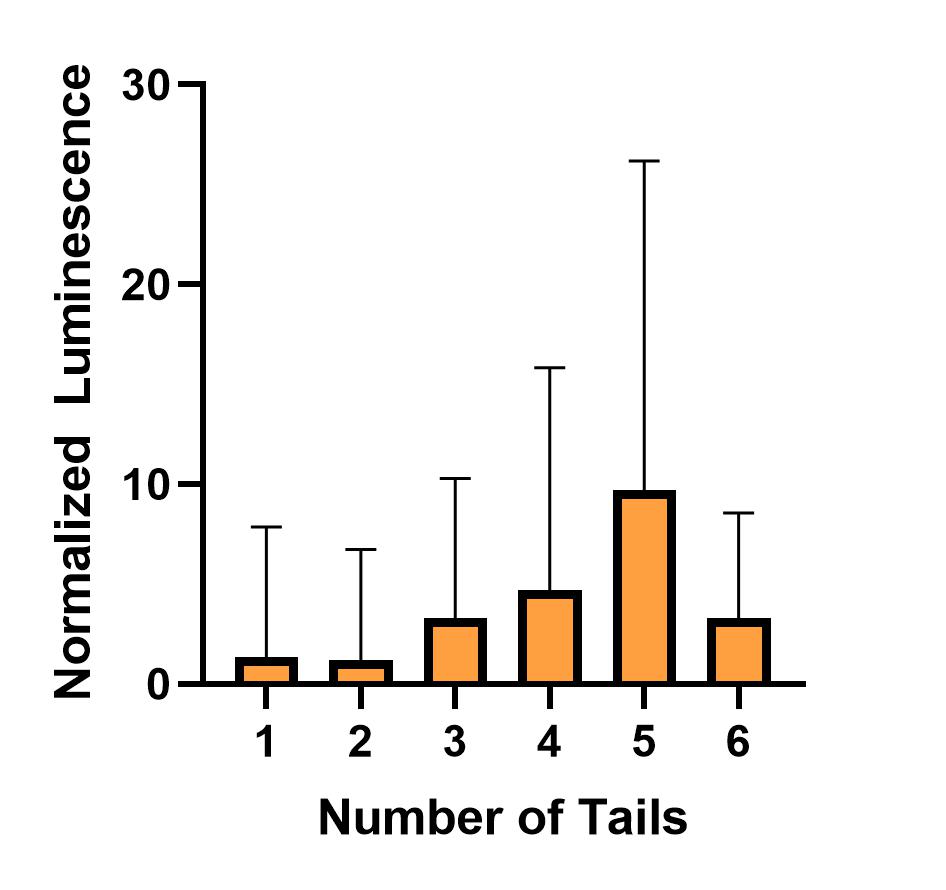d | 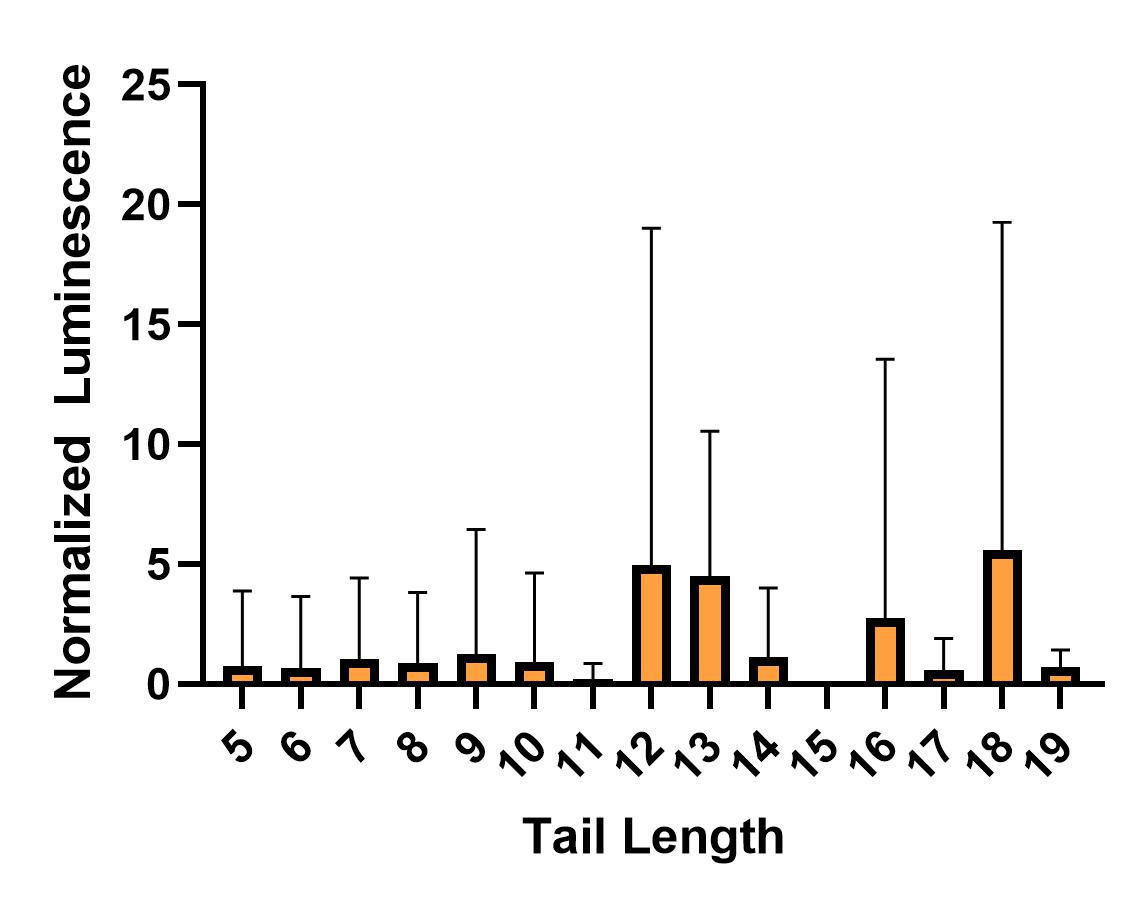e |  |

**Figure S1.** **Distribution of luminescence within groups of ionizable lipid features.** (a) Number of amines. (b) Number of esters. (c) Number of rings. (d) Number of tails. (e) Tail length.

**Extended Methods**

**Database Creation:** Luminescence values for each LNP are extracted from heatmaps by loading the image into PowerPoint and using the eyedropper tool to attain the RGB code for each point. A linear equation for the highest and lowest value on each heatmap is created using this method. Finally, the linear equation is used to calculate the corresponding luminescence values from the RGB codes. Luminescence values are extracted from bar graphs by measuring the height of each bar with a ruler.

**Normalization and Curation of the Database:** Each paper is normalized using an experimentally tested or calculated using either Onpattro, C12-200 DSPC, or lipofectamine control. For studies evaluating more than one dose, the highest dose is input into the database. Unless establishment of an alternate control is described below, the authors directly tested Onpattro.

Combinatorial design of nanoparticles for pulmonary mRNA delivery and genome editing^1^

An array of 72 head groups, 10 tail groups, and 1 linker are synthesized via 3-component reaction chemistry to form ionizable lipids to formulate LNPs for testing in A549s. Subsequent testing through intramuscular injection leads to the identification of the lead candidate, RCB 4-8. Although there are no internal controls for the in vitro experiment, a second in vivo study through intratracheal delivery compares the lead candidate to Onpattro. Therefore, we establish an in vitro in vivo correlation factor to correlate the performance on Onpattro in vivo to its performance in vitro in A549s.

To establish this value, the luminescence value of the highest intratracheal dosing of lead candidate RCB 4-8 (0.5 mg/kg) and divided by the luminescence of the in vitro lead candidate (A11-T1). As a note, the intratracheal value for RCB 4-8 was not divided by its corresponding in vitro value because there is no correlation between the two luminescence values – the inferior in vitro efficacy of RCB 4-8 was not reflective of its superior performance through intramuscular delivery. The Onpattro control was created by dividing the intratracheal dosing of Onpattro by the resulting in vitro in vivo correlation factor.

In vitro in vivo correlation factor:

3.475x10^8^ / 9.37x10^5^ = 371

Raw value of Onpattro control:

4.140x10^6^ / 371 = 1.12x10^4^

A Combinatorial Library of Lipid Nanoparticles for Cell Type-Specific mRNA Delivery^2^

A total of 10 lipids containing a tertiary amine head group, two levels of linker backbone (hydroxylamine or ethanolamine), and four types of hydrophobic tails are synthesized, and LNPs encapsulating firefly luciferase mRNA are tested in three different cell types (B16F10, CT26, and Raw264.7).

Lipid 6 and Lipid 8 share a high level of structural similarity to MC3: they both have linoleoyl fatty acid chains as tails and a primary tertiary amine. Additionally, Lipid 6 and Lipid 8 behave similarly when formulated in an LNP encapsulating siRNA for PLK1 (a marker with a role in cell viability) compared to an MC3 LNP^3^. As Lipid 6 is the most structurally similar to MC3 compared to Lipid 8, it serves as the internal control.

Lipid Nanoparticle Composition Drives mRNA Delivery to the Placenta^4^

A DOE approach is used to design formulations containing C12-200 or MC3 as the ionizable lipid and DOPE or DSPC as the helper lipid. The ratios of the ionizable lipid range from 25-45%, the helper lipid from 10-22%, and the PEG-Lipid from 1.5-3.5%. While the study does not formulate Onpattro directly, the formulation parameters of A7 (MC3:DSPC:DMPE-PEG:Cholesterol 45:10:1.5:43.5) are the most similar to Onpattro’s ratios (50:10:1.5:38.5). As all LNPs formulated with MC3 and DSPC across different ratios in this paper have similar luciferase activity compared to the lead candidate, A7 serves as an internal control.

Aerosolizable Lipid Nanoparticles for Pulmonary Delivery of mRNA through Design of Experiments^5^

A DOE approach is used to design formulations containing the ionizable lipid MC3 and using the helper lipids DOPE, DSPC, and DPPC and using the PEG-Lipids DMPE-PEG, DMG-PEG, and DSPE-PEG. The ratios of the ionizable lipid range from 40-60%, the helper lipid from 10-20%, the PEG-Lipid from 1-5%, and the cholesterol from 15-49%. Although Onpattro is not directly tested, the closest formulation is F17 (MC3:DOPE:DMG-PEG:Cholesterol 40:16:1:43). A key basis for this assumption is that LNPs formulated in this paper performed similarly at 1% PEG across helper lipids – LNPs formulated with 3 or 5% PEG performed significantly worse.

Optimization of Lipid Nanoparticles for saRNA Expression and Cellular Activation Using a Design-of-Experiment Approach^6^

A DOE approach is used to design LNPs encapsulating luciferase saRNA formulated with three levels of ionizable lipid (MC3, SM-102, or ALC-0315), two levels of helper lipid (DOPE or DSPC), three levels of formulation buffer pH (4, 5, 6), and an ionizable lipid range from 35-45%, a helper lipid range from 12-17.5%, and a constant 1.25% PEG-Lipid ratio. In this case, the closest formulation to Onpattro is LNP 3 (MC3:DOPE:DMG-PEG:Cholesterol 40:15:1.25:43.75). As a note, only LNPs formulated with pH 4 are considered. Raw values from this study are kindly provided by Dr. Blakney.

Delivery of mRNA vaccines with heterocyclic lipids increases anti-tumor efficacy by STING-mediated immune cell activation^7^

An ionizable lipid library of 1,080 lipids with 12 primary or secondary amines, 10 isocyanide derivatives, 9 alkyl ketones is synthesized through isocyanide-mediated 3 component reaction chemistry. The LNPs encapsulating firefly luciferase mRNA were tested in HeLa cells (Figure 2a). The lead candidate, A252DC18, is chosen from testing in HeLa, BMDC, and BMDM cell lines. Although MC3 is not tested in these studies, it is tested in a later in vivo study. Mice were vaccinated with LNPs encapsulating the mRNA encoding for the OVA peptide, and the IFN-γ expression is measured after restimulation of the splenocytes with the peptide ex vivo. In this study, the lead candidate in vitro performed similarly with MC3. Therefore, A252DC18 was selected as the internal control. Additionally, there is another study in the supplementary (Table S1) that tests the effect of lipid/mRNA weight ratio and different ratios of A2, DOPE, and PEG of the luciferase activity of the lead candidate. This supplementary study and the first study are normalized through F8 (A2:DOPE:C14-PEG:Cholesterol 50:16:1.5:32.5), which has the closest formulation to Onpattro.

Ionizable Lipid Nanoparticle-Mediated mRNA Delivery for Human CAR T Cell Engineering^8^

Ionizable lipids are synthesized via Michael addition of 3 alkyl chains and 8 polyamine cores for a final library of 24 lipids. LNPs are tested in Jurkat cells and compared with lipofectamine. As Ly et al. also uses lipofectamine, a correlation factor between lipofectamine and this study’s Onpattro control is calculated. To create the internal Onpattro control, the value of lipofectamine from Billingsley et al. is divided by this Lipofectamine-Onpattro correlation factor.

Lipofectamine from Ly et al. = 1.56x10^6^

LNP 3 from Ly et al. = 5.74x10^5^

Lipofectamine-Onpattro correlation factor: Lipofectamine / LNP 3 = 2.72

Lipofectamine from Billingsley et al. = 0.2

Lipofectamine from Billingsley et al./ Lipofectamine-Onpattro correlation factor = 0.2 / 2.72 = 0.074

Imidazole-Based Synthetic Lipidoids for In Vivo mRNA Delivery into Primary T Lymphocytes^9^

A library of cationic lipid-like materials is synthesized through Michael addition of 20 hydrophilic amine head and 13 hydrophobic carbon tails. In this study, not all possible reactions are carried to completion. Therefore, the final number of possible synthesis reactions (260) is not encompassed in the final number (78) of LNPs tested in primary human CD8+ T lymphocytes. These LNPs are compared against lipofectamine, and a Lipofectamine-Onpattro correlation factor (2.72) is used to establish an Onpattro control as previously discussed for Billingsley et al.

Lipofectamine from Zhao e al. / Lipofectamine-Onpattro correlation factor = 6.50x10^3^/2.72 = 2.4x10^3^

A second in vitro study is conducted using analogs of the 93 head group from the first study that showed superior delivery when synthesized with the 3 tail variants containing 17 carbons. They test 18 new head groups with the same 3 tail variants to total 54 analogs. As these new LNPs were compared with the LNPs formulated in the first study with the 93 head group, a first study – second study factor is established with the lead candidate (93-O17O). The raw value from each of the LNPs in the second study is then multiplied by this factor. Once the raw values between studies are normalized, the Lipofectamine-Onpattro correlation factor is similarly applied to the values in the second study. This process is repeated for a third study in the supplementary testing different ratios of an LNP formulated with the 93-O17S (Figure S2a).

93-O17O from first study = 1.22x10^4^

93-O17O from second study = 9.94x10^3^

First study – Second study factor for 93-O17O = 1.22x10^4^/9.94x10^3^ = 1.23

93-O17S from first study = 1.13x10^4^

93-O17S from third study = 4.32x10^3^

First study – Third study factor for 93-O17S = 1.13x10^4^/4.32x10^3^ = 2.62

An Orthogonal Array Optimization of Lipid-like Nanoparticles for mRNA Delivery in Vivo^10^

Ionizable lipids generated from multi-step chemistry from N1,N3,N5-tris(2-aminoethyl)benzene-1,3,5-tricarboxamide (TT) derivatives. These derivates are tested in Hep3B cells, and a lead candidate (TT3) is chosen for subsequent analysis. Two rounds of DOE experiments are then performed using varying levels of DOPE, Cholesterol, and DMG-PEG for a total of 32 formulations. Although none of the LNPs in the three studies are the same, the second round of DOE compared the transfection efficiency with a C12-200 DSPC LNP. Based on previous work^11^, C12-200 LNPs formulated with DSPC helper lipid perform similarly. As a study used in this analysis^4^ formulates an LNP with C12-200 and DSPC (A15: C12-200:DSPC:DMG-PEG:Cholesterol 45:22:1.5:31.5), this LNP thus serves as the internal control. The raw value of all LNPs in this study is divided by the raw value of A15.

**References**

1.	Li B, Manan RS, Liang SQ, Gordon A, Jiang A, Varley A, Gao G, Langer R, Xue W, Anderson D. Combinatorial design of nanoparticles for pulmonary mRNA delivery and genome editing. *Nat Biotechnol*. Published online March 30, 2023:1-6. doi:10.1038/s41587-023-01679-x

2.	Naidu GS, Yong SB, Ramishetti S, Rampado R, Sharma P, Ezra A, Goldsmith M, Hazan-Halevy I, Chatterjee S, Aitha A, Peer D. A combinatorial library of lipid nanoparticles for cell type-specific mRNA delivery. *Adv Sci*. 2023;10(19):2301929. doi:10.1002/advs.202301929

3.	Ramishetti S, Hazan-Halevy I, Palakuri R, Chatterjee S, Gonna SN, Dammes N, Freilich I, Shmuel LK, Danino D, Peer D. A combinatorial library of lipid nanoparticles for RNA delivery to leukocytes. *Adv Mater*. 2020;32(12):1906128. doi:10.1002/adma.201906128

4.	Young RE, Nelson KM, Hofbauer SI, Vijayakumar T, Alameh MG, Weissman D, Papachristou C, Gleghorn JP, Riley RS. Lipid nanoparticle composition drives mRNA delivery to the placenta. Published online December 22, 2022:2022.12.22.521490. doi:10.1101/2022.12.22.521490

5.	Zhang H, Leal J, Soto MR, Smyth HDC, Ghosh D. Aerosolizable lipid nanoparticles for pulmonary delivery of mRNA through design of experiments. *Pharmaceutics*. 2020;12(11):1042. doi:10.3390/pharmaceutics12111042

6.	Ly HH, Daniel S, Soriano SKV, Kis Z, Blakney AK. Optimization of lipid nanoparticles for saRNA expression and cellular activation using a design-of-experiment approach. *Mol Pharmaceutics*. 2022;19(6):1892-1905. doi:10.1021/acs.molpharmaceut.2c00032

7.	Miao L, Li L, Huang Y, Delcassian D, Chahal J, Han J, Shi Y, Sadtler K, Gao W, Lin J, Doloff JC, Langer R, Anderson DG. Delivery of mRNA vaccines with heterocyclic lipids increases anti-tumor efficacy by STING-mediated immune cell activation. *Nat Biotechnol*. 2019;37(10):1174-1185. doi:10.1038/s41587-019-0247-3

8.	Billingsley MM, Singh N, Ravikumar P, Zhang R, June CH, Mitchell MJ. Ionizable lipid nanoparticle-mediated mRNA delivery for human CAR T cell engineering. *Nano Lett*. 2020;20(3):1578-1589. doi:10.1021/acs.nanolett.9b04246

9.	Zhao X, Chen J, Qiu M, Li Y, Glass Z, Xu Q. Imidazole-based synthetic lipidoids for in vivo mRNA delivery into primary T lymphocytes. *Angew Chem, Int Ed Engl*. 2020;59(45):20083-20089. doi:10.1002/anie.202008082

10. Li B, Luo X, Deng B, Wang J, McComb DW, Shi Y, Gaensler KML, Tan X, Dunn AL, Kerlin BA, Dong Y. An orthogonal array optimization of lipid-like nanoparticles for mRNA delivery in vivo. *Nano Lett*. 2015;15(12):8099-8107. doi:10.1021/acs.nanolett.5b03528

11.	Kauffman KJ, Dorkin JR, Yang JH, Heartlein MW, DeRosa F, Mir FF, Fenton OS, Anderson DG. Optimization of lipid nanoparticle formulations for mRNA delivery in vivo with fractional factorial and definitive screening designs. *Nano Lett*. 2015;15(11):7300-7306. doi:10.1021/acs.nanolett.5b02497
